## Supplementary Information for "Dynamics of activation in the voltage-sensing domain of Ci-VSP"

### S1. SUPPLEMENTARY METHODS

#### A. Mean first passage time

Like the committor, the mean first passage time (MFPT) from a point $x$ to a state $B$ also satisfies a boundary value equation involving the stopped transition operator (that stops trajectories when they enter $B$):

$$m(x) = \begin{cases} \mathcal{S}_t m(x) = -1 & x \in B^c \\ 0 & x \in B \end{cases} \tag{S1}$$

For the MFPT calculations, we use an indicator basis with modified features. We construct the basis by computing the top ten eigenvectors from the 60 input features used for the committor with IVAC and lag times between 1 and 100 ns. The resulting 10-dimensional

*) Electronic mail:

†) Electronic mail:

data is clustered with $k$-means ($k = 200$), and the resulting clusters are treated as sets that define indicator basis functions. We also modify the state definition of $B$ to

$$\left(\frac{\max\{d + 4.24\ \text{Å}, 0\ \text{Å}\}}{1.1\ \text{Å}}\right)^2 + \left(\frac{\max\{\theta + 56.95^\circ, 0^\circ\}}{8^\circ}\right)^2 < 1$$

so that states beyond the up state (including the up+ state) do not contribute to the MFPT. The salt-bridge cutoffs (Table 1) are maintained inside the original ellipse defining $B$. Finally, we employ a memory-based algorithm that empirically improves the stability and accuracy of the results[1–3].

#### B. Backward committor

Analogously to the forward committor, the backward committor $q_-$ represents the probability of having last exited from $A$, the reactant state, rather than $B$[4]. The backward committor also satisfies a boundary value equation involving the stopped transition operator:

$$q_-(x) = \begin{cases} \mathcal{S}_{-t} q_-(x) & x \in (A \cup B)^c \\ 1 & x \in A \\ 0 & x \in B \end{cases} \tag{S2}$$

where $\mathcal{S}_{-t}$ is the backwards-in-time transition operator stopped upon first entrance into $D^c$. The basis set used for calculation of $q_-$ is the same pairwise-distance basis as used for the forward committor, though the guess function is amended to $\gamma(x) = d_B^2/(d_A^2 + d_B)^2$. For a reversible system (i.e., one that satisfies detailed balance, as does Ci-VSP at 0 mV), the backward committor should satisfy $q_- = 1 - q_+$, though this does not hold exactly in practice due to statistical errors in estimating committors.

When plotting committors, we use $\mathbf{E}_\pi[\mathbf{1}_{\{\xi \in ds\}}(x) \mathcal{S}_\tau q_+(x)]$, where $\mathbf{1}_{\{\xi \in ds\}}(x)$ is an indicator function on a bin in our chosen CV space, e.g., $\xi(x) = (d(x), \theta(x))$, and $\pi$ is the stationary distribution, obtained by reweighting our sampling distribution $\mu$. The propagation via $\mathcal{S}_\tau$ alleviates some of the projection error incurred in computing $q_+$ using DGA.

### S2. SUPPLEMENTARY TABLES

Supplementary Table 1. Salt bridge distance cutoffs for down and up states

| **Salt bridge** | **Cutoff (Å)** |
|---|---|
| *Down* | |
| R226 $C_\zeta$–D129 $C_\delta$ | $> 6.0$ |
| R226 $C_\zeta$–D186 $C_\delta$ | $< 5.0$ |
| *Up* | |
| R226 $C_\zeta$–D129 $C_\delta$ | $> 5.0$ |
| R229 $C_\zeta$–D129 $C_\delta$ | $< 11.0$ |
| R229 $C_\zeta$–D186 $C_\delta$ | $< 7.5$ |
| R232 $C_\zeta$–D186 $C_\delta$ | $< 6.0$ |

Supplementary Table 2. Collective variables considered in this work

| **Type** | **Name** | **Description** |
|---|---|---|
| S4 helix | translocation | translocation along principal axis |
| | rotation | rotation around principal axis |
| salt bridge | R226 $C_\zeta$-D129 $C_\gamma$ | salt-bridge distances between Arg and Asp/Glu side chains |
| | R226 $C_\gamma$-D186 $C_\delta$ | |
| | R229 $C_\zeta$-E183 $C_\delta$ | |
| | R229 $C_\zeta$-D186 $C_\gamma$ | |
| | R232 $C_\zeta$-E183 $C_\delta$ | |
| | R232 $C_\zeta$-D186 $C_\gamma$ | |
| hydrogen bonding | R226-D129 | hydrogen bonds between salt-bridge residue side chains (sensing arginines and countercharges) |
| | R226-D186 | |
| | R229-E183 | |
| | R229-D186 | |
| | R232-E183 | |
| | R232-D186 | |
| hydrophobic plug | R226-F161 | distances between Arg $C_\zeta$ and center-of-mass of Phe phenyl group |
| lipid headgroup | R217 | hydrogen bonds between arginine guanidinium and phosphate groups of POPC lipids |
| | R229 | |
| | R232 | |
| arginine hydration | R223 | hydrogen bonds between arginine side chain and water |
| | R229 | |
| | R229 | |
| | R232 | |

### S3. SUPPLEMENTARY FIGURES

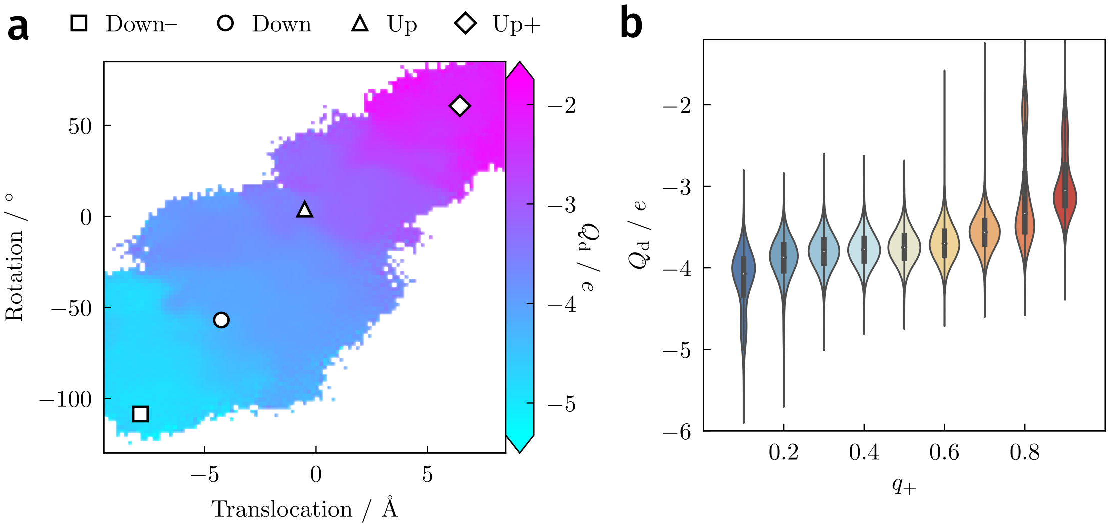


Supplementary Figure 1. Displacement charge. **a**, Average $Q_\mathrm{d}$ for the full system plotted against the S4 helix CVs. **b**, Distribution of displacement charge as a function of the committor. Box plots indicate median and interquartile range for each committor range.

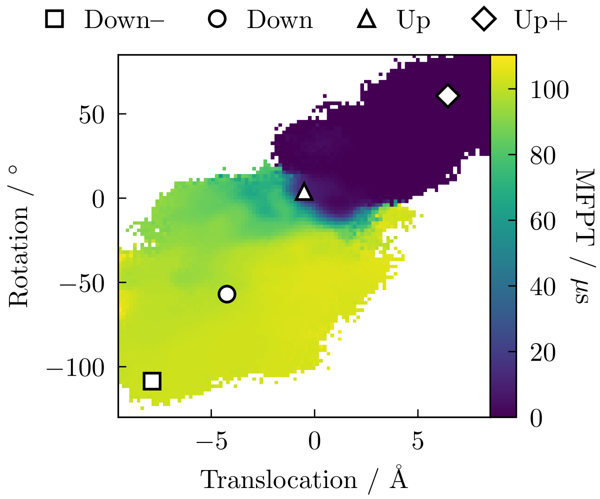


Supplementary Figure 2. Mean first-passage time (MFPT) to the up state as a function of the S4 helix CVs. The average MFPT for points in the down state is approximately 100 $\mu$s.

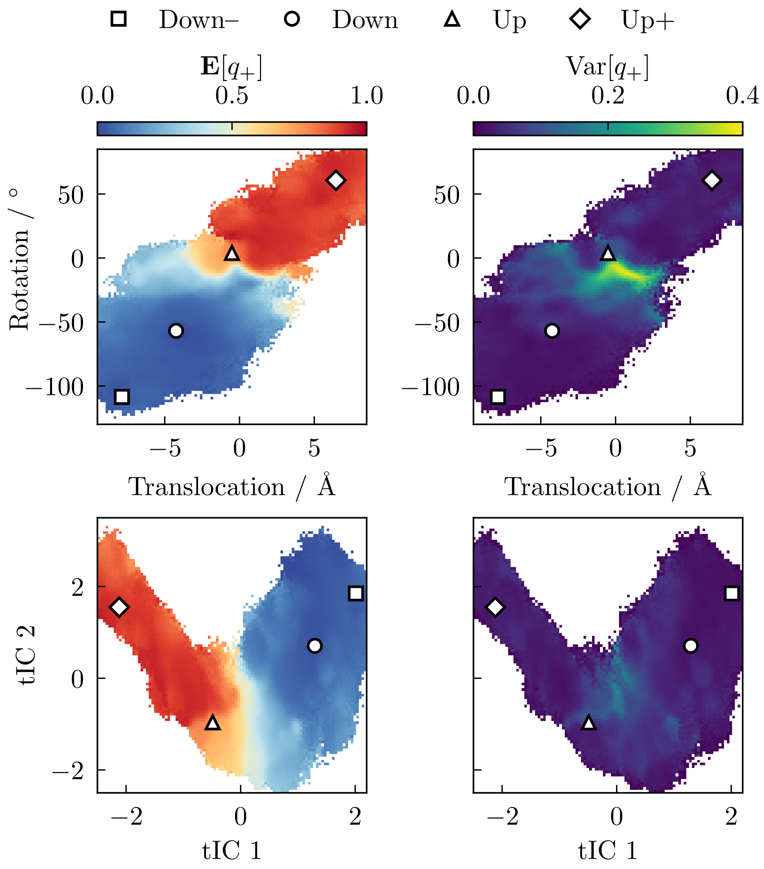


Supplementary Figure 3. Variance in committor estimates for two different CV projections. Top, S4 helix CVs. Bottom, first two IVAC coordinates. Right, the variance is computed and normalized for each CV histogram bin. Left, the average committor is reproduced from the main text for visual reference.

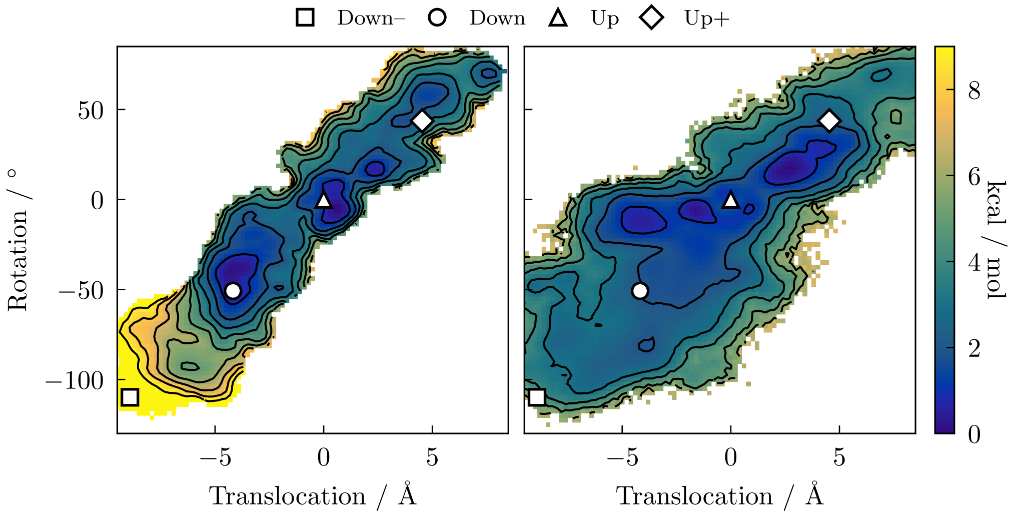


Supplementary Figure 4. Comparison of PMF along S4 helix coordinates computed by REUS[5] (left) and DGA (right).

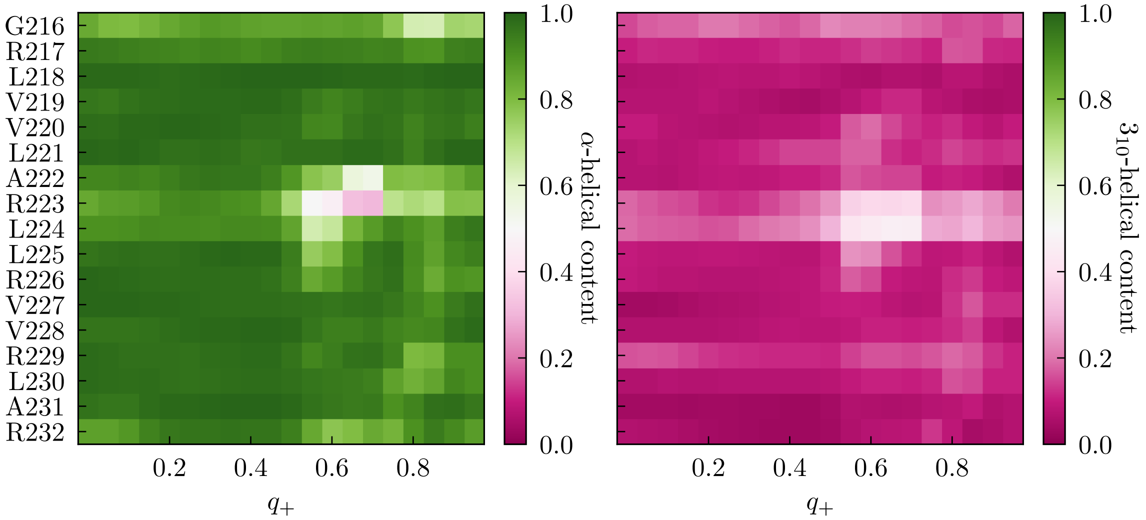


Supplementary Figure 5. $\alpha$-helical (left) or $3_{10}$-helical (right) content of S4 residues as a function of $q_+$. Residue $i$ is classified as adopting an $\alpha$-helical or a $3_{10}$-helical conformation if it forms a hydrogen bond ($d_{\mathrm{O-N}} < 3.5$ Å and $\angle_{\mathrm{O-H-N}} > 120°$) to residue $i+4$ or $i+3$, respectively[5].

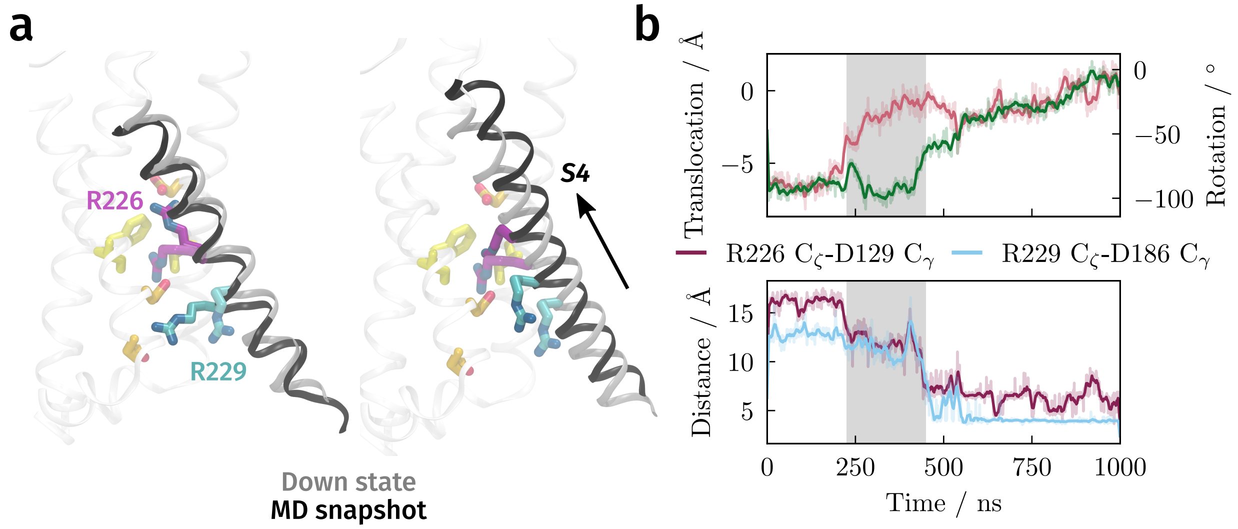


Supplementary Figure 6. **a**, Selected frames from a simulation (opaque) superimposed against the down state crystal structure (translucent) after aligning the S1–S3 helices. Only the S4 helix, R226, and R229 of the MD simulation frame are shown for clarity. **b**, Top, translocation (pink) and rotation (green) of S4 helix. Bottom, salt-bridge distances during a representative 1 $\mu$s trajectory showing translocation of the helix preceding of rotation and movement of the arginine side chains. Light lines show instantaneous values, while dark lines show moving averages over ten frames.

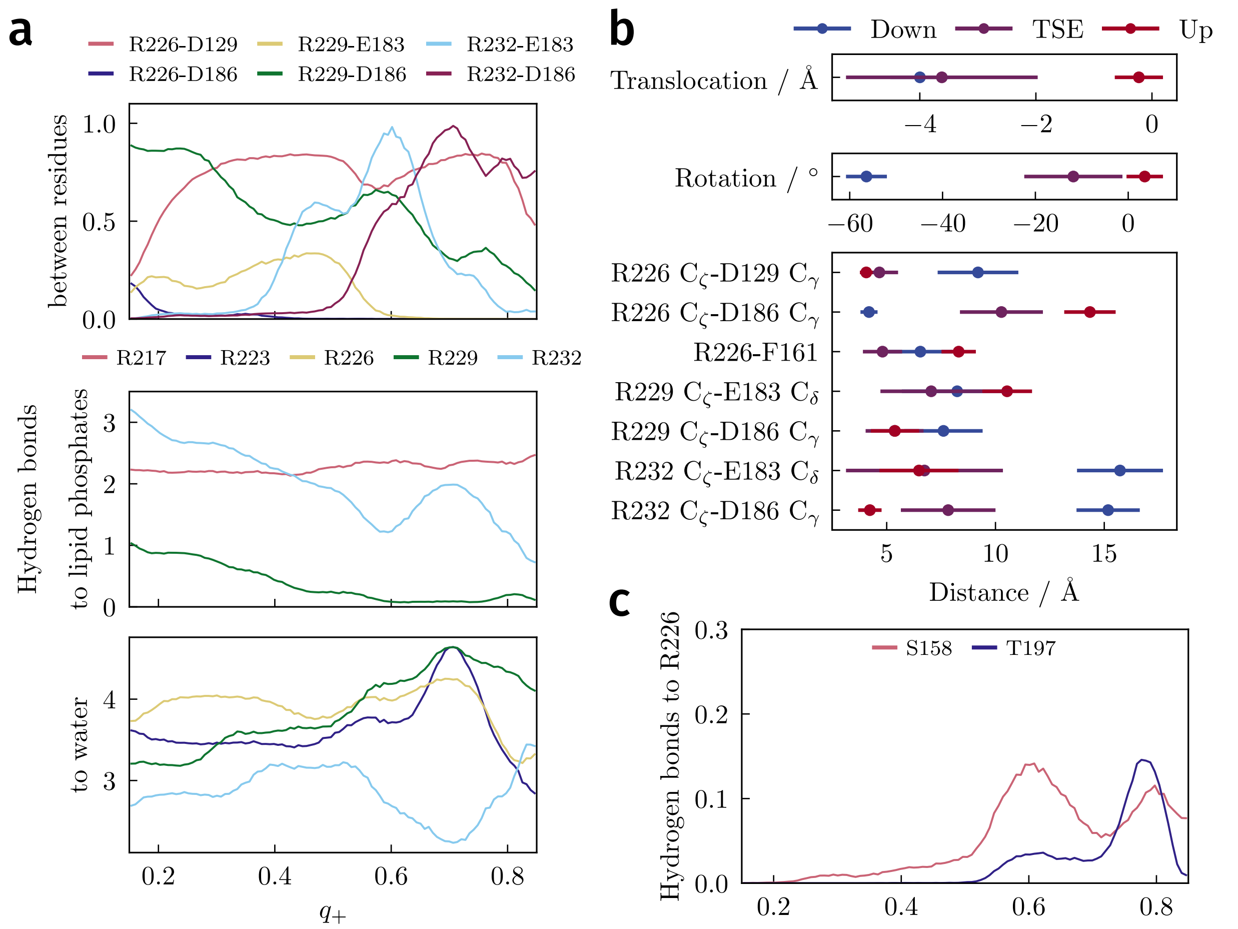


Supplementary Figure 7. Transition state ensemble and hydrogen bonding. **a**, Average number of hydrogen bonds between arginines and acidic amino acids (top), to lipid headgroups (center), and to water (bottom). Instantaneous hydrogen bond values are obtained by averaging over a moving window of 10 ns (100 frames). **b**, Average of selected CVs in the down state, in the transition state ensemble ($q_+ \in [0.45, 0.55]$), and in the up state. **c**, Average number of hydrogen bonds between R226 and S158 or T197.

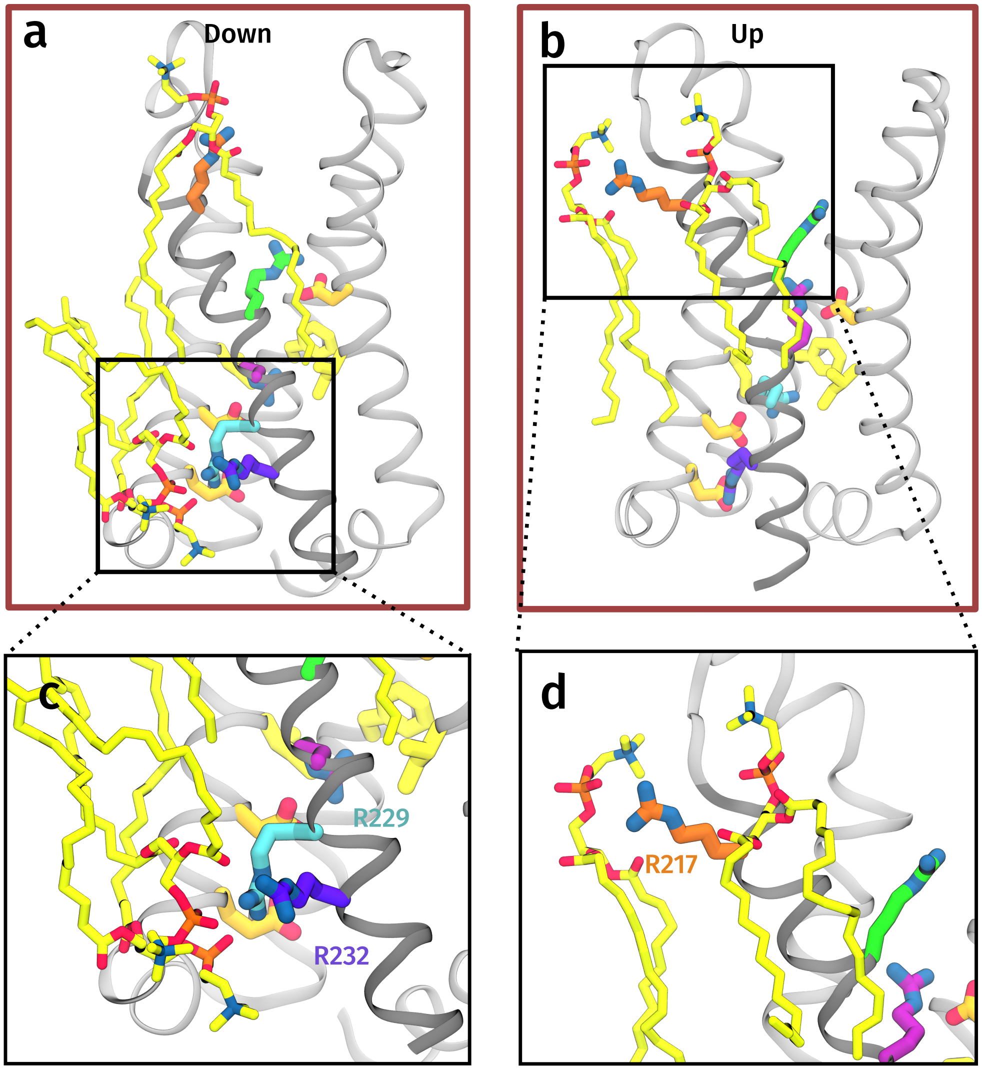


Supplementary Figure 8. Interactions between sensing arginines and lipids. **a** and **b**, Down (**a**) and up (**b**) states with nearby lipid molecules drawn. **c** and **d**, Interactions between R229/R232 in the down state (**c**) or R217 in the up state (**d**) with headgroups of POPC lipids. Phosphates and oxygens in the phosphatidylcholine headgroup are colored in orange and red, respectively, whereas carbons of lipid tails are in yellow.

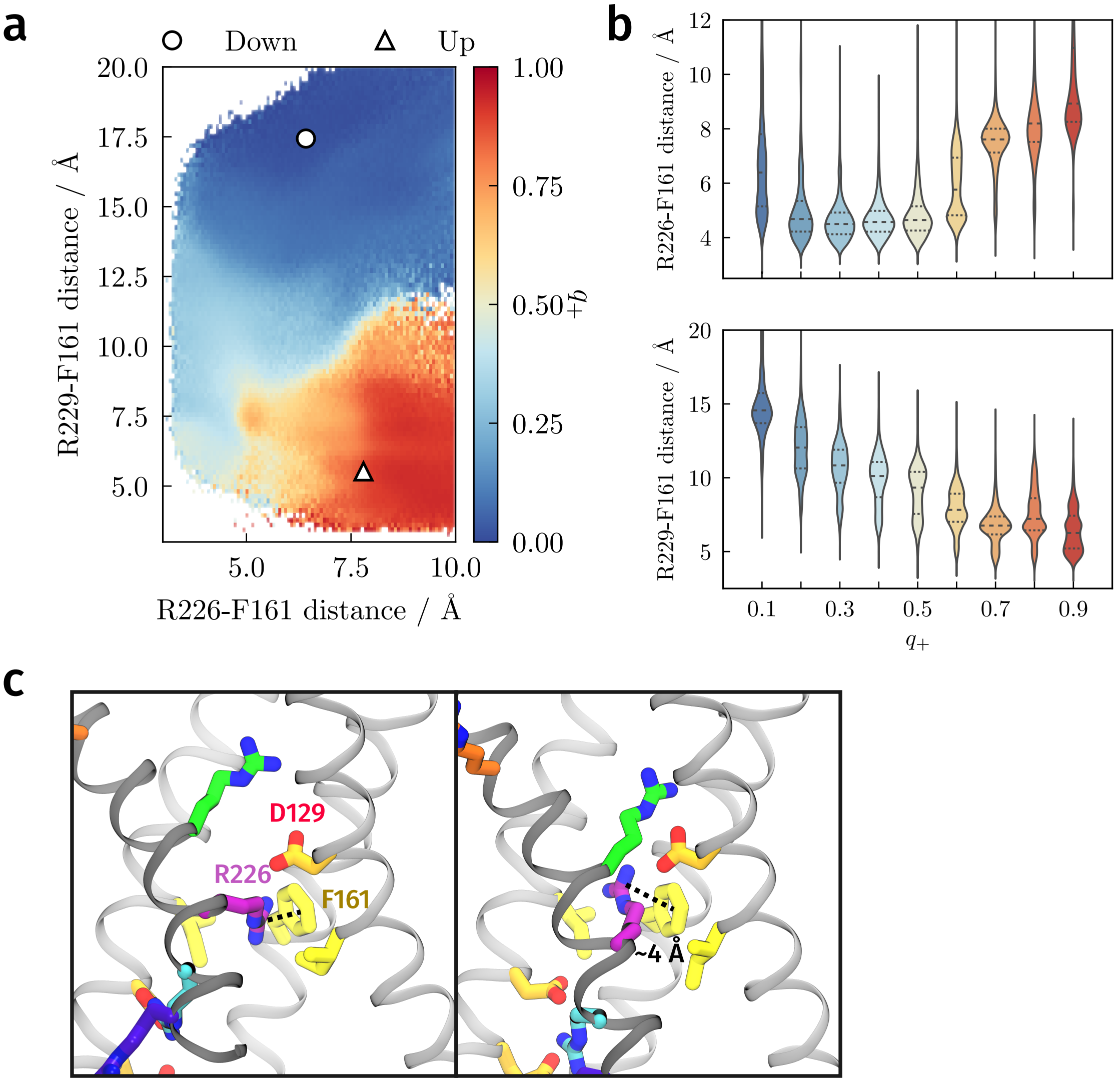


Supplementary Figure 9. Interactions between R226, R229, and F161. **a**, Committor projected onto R226-F161 and R229-F161 distances. **b**, Distribution of R226-F161 and R229-F161 distances as a function of the committor. Each violin is obtained by binning structures with the indicated $q_+ \pm 0.05$ (e.g., the first violin contains structures with $q_+ \in [0.15, 0.25]$). The median and upper/lower quartiles are denoted with dashed and solid lines, respectively. **c**, Representative configurations of R226, F161, and nearby residues.

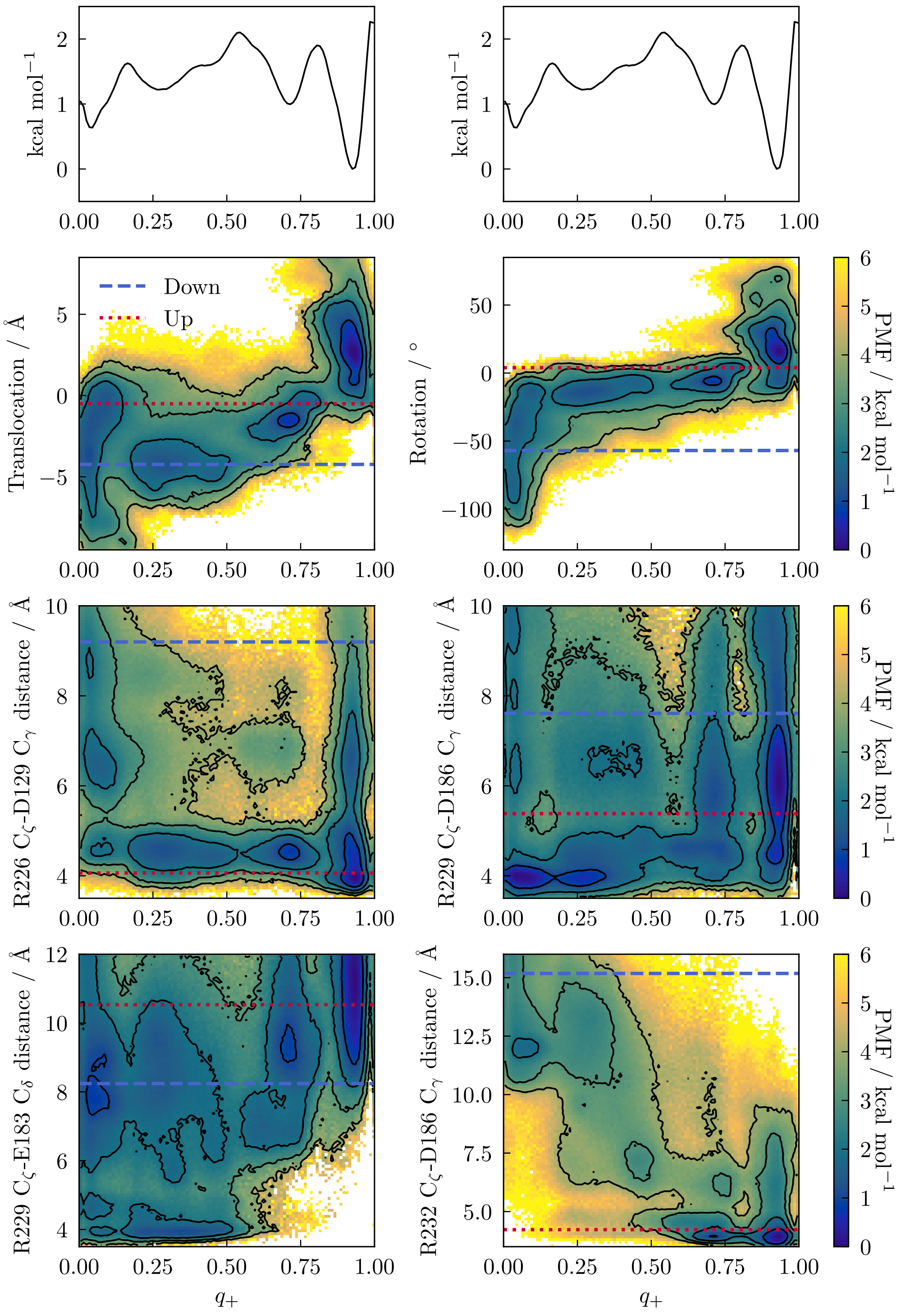


Supplementary Figure 10. PMF projected onto committor and selected CVs (describing the S4 helix and salt-bridges) with the projection of the PMF along the committor above. Contours on the two-dimensional PMFs are drawn every 1 kcal/mol.

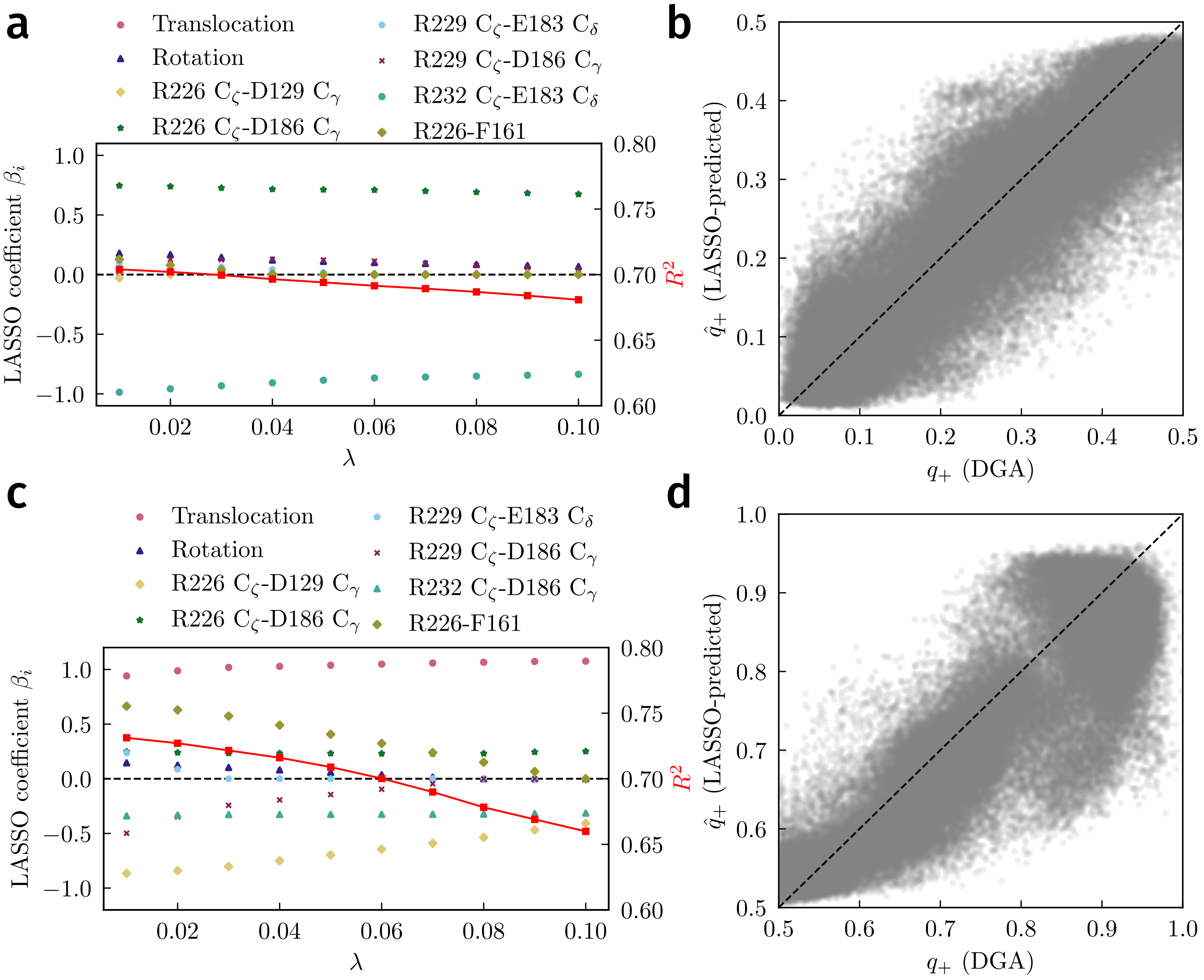


Supplementary Figure 11. Sparse regression of the committor. **a**, Changes in coefficients from LASSO fitting for the points down → TS as a function of regularization strength. Only nonzero coefficients from $\lambda = 0.02$ are shown. The goodness-of-fit, $R^2$ is plotted as red squares. **b**, Scatter plot of DGA committors versus LASSO committors. **c**, Changes in coefficients from LASSO fitting for the points TS → up as a function of regularization strength. Only nonzero coefficients from $\lambda = 0.03$ are shown. The goodness-of-fit, $R^2$ is plotted as red squares. **d**, Scatter plot of DGA committors versus LASSO committors.

### S4. SUPPLEMENTARY MOVIES

Supplementary Movie 1. Down-up transition in Ci-VSD. Five configurations are sampled from twenty equally spaced windows of the committor (i.e., of width 0.05) and are aligned. The red line indicates the progress of the transition along the committor.

### SUPPLEMENTARY REFERENCES

[1] J. Strahan, S. C. Guo, C. Lorpaiboon, A. R. Dinner, and J. Weare, "Inexact iterative numerical linear algebra for neural network-based spectral estimation and rare-event prediction," *arXiv:2303.12534 [physics.comp-ph]* (2023).

[2] S. Cao, A. Montoya-Castillo, W. Wang, T. E. Markland, and X. Huang, "On the advantages of exploiting memory in Markov State Models for biomolecular dynamics," *J. Chem. Phys.* **153**, 014105 (2020).

[3] E. Darve, J. Solomon, and A. Kia, "Computing generalized Langevin equations and generalized Fokker–Planck equations," *Proc. Natl Acad. Sci. USA* **106**, 10884–10889 (2009).

[4] E. Vanden-Eijnden, "Transition path theory," in *Computer Simulations in Condensed Matter Systems: From Materials to Chemical Biology Volume 1* (Springer, 2006) pp. 453–493.

[5] R. Shen, Y. Meng, B. Roux, and E. Perozo, "Mechanism of voltage gating in the voltage-sensing phosphatase Ci-VSP," *Proc. Natl Acad. Sci. USA* **119**, e2206649119 (2022).
